# A systems-level model of sleep-dependent memory-consolidation failure in neurodegeneration: the spindle–slow-oscillation decoupling cascade dissociates amyloid and tau

**DOI:** 10.64898/2026.09.21.753161

**Authors:** Bahaa Masry, Anas Shahin, Zeina S. Malek

## Abstract

During non-rapid-eye-movement (NREM) sleep, the temporal coupling of cortical slow oscillations (SOs), thalamic spindles, and hippocampal sharp-wave ripples drives the consolidation of declarative memories. This coupling degrades in ageing and Alzheimer’s disease (AD), and although A*β* and tau leave dissociable signatures in human sleep, the mechanisms by which progressive pathology dismantles the consolidation machinery are difficult to isolate experimentally, and have not to our knowledge been reproduced in a model that can be perturbed directly. We built a systems-level model in which cortical SOs and thalamic spindles are generated by reduced oscillators, hippocampal ripples replay encoded spike sequences, and the *measured* per-event SO–spindle timing alignment causally gates spike-timing-dependent plasticity on cortical sequence synapses. A post-sleep cued-recall test reads out consolidation. Five neurodegeneration parameters (amyloid, tau, synaptic density, GABAergic inhibition, cholinergic tone) map to distinct mechanisms grounded in the human and animal literature. The model reproduces graded healthy consolidation and a progressive collapse in which coupling, slow-wave power, spindle power and recall fall monotonically and the overnight memory effect flips from consolidation to net forgetting, with weak memories failing first. Scrambling SO–spindle timing while holding oscillation power fixed abolishes consolidation, establishing that coupling timing, rather than oscillation power, is what the plasticity gate depends on within the model. A*β* and tau impair memory through orthogonal signatures (A*β* collapses slow-wave power while sparing replay order, tau the reverse) and this orthogonality holds across the entire A*β*× tau plane and survives simultaneous ±50% resampling of every mapping coefficient (40/40 samples), so it is not an artefact of a single calibration point. The model yields a falsifiable clinical prediction: closed-loop slow-oscillation enhancement rescues memory only when the deficit is amplitude/coupling-dominated, not when it is replay(tau)-dominated, despite normalising slow-wave power in both cases. Because the therapy arms dissociate coupling from memory benefit, the model also cautions against adopting SO–spindle coupling as a standalone surrogate endpoint.

**Author summary:** Sleep strengthens new memories through a precisely timed dialogue between three brain rhythms: cortical slow waves, thalamic sleep spindles, and hippocampal ripples. This timing breaks down in Alzheimer’s disease, but it has been unclear how the two hallmark pathologies of the disease (amyloid plaques and tau tangles) each contribute, and whether the same treatment would help both. We built a computer model that reproduces the rhythms and, crucially, lets the *measured* timing between slow waves and spindles control how strongly memories are written into cortex. As we gradually increase simulated pathology, the rhythms uncouple, memory consolidation fails, and overnight the model begins to *forget* rather than remember, exactly the trajectory seen in patients. When we scramble only the timing of the rhythms while leaving their strength untouched, consolidation collapses, showing that timing itself carries the memory benefit. The model separates amyloid (which weakens slow waves) from tau (which corrupts the hippocampal replay that timing is supposed to gate), and predicts that a leading experimental therapy (slow-wave stimulation during sleep) will help amyloid-dominated patients but fail tau-dominated patients even though it restores their slow waves. This prediction is directly testable and could guide patient selection for sleep-based interventions.

## Introduction

Sleep is not a passive state for memory. The active systems consolidation framework holds that, during NREM sleep, newly encoded hippocampal traces are repeatedly reactivated and gradually redistributed to neocortex for long-term storage [1, 2]. This redistribution is orchestrated by the precise temporal nesting of three cardinal oscillations: the cortical slow oscillation (SO, ~0.5–1 Hz), the thalamocortical sleep spindle (~10–16 Hz), and the hippocampal sharp-wave ripple (~150–250 Hz). Complementary learning systems theory provides the computational rationale: a fast hippocampal learner instructs a slow cortical learner offline to avoid catastrophic interference [3]. Intracranial recordings in humans show that these rhythms are hierarchically coupled, with spindles grouped on the SO up-state and ripples nested within spindle troughs [4, 5], and causal optogenetic work in rodents shows that only spindles delivered *in phase* with the SO up-state enhance hippocampus-dependent memory [6]. The strength of SO–spindle phase–amplitude coupling predicts overnight memory across the lifespan [7, 8].

This coupling deteriorates in ageing and AD, and the deterioration tracks the two molecular hallmarks of the disease in dissociable ways. Medial-prefrontal A*β* burden impairs the generation of NREM slow-wave activity (SWA), and this SWA deficit statistically mediates impaired hippocampus-dependent memory [9]. In contrast, medial-temporal tau burden is preferentially associated with the loss of SO–spindle coupling and with reduced NREM slow-wave activity at 1–2 Hz [10, 11], and pathological tau slows the slow oscillation and degrades hippocampal ripple-associated spike dynamics in mouse models [12, 13]. Reduced sharp-wave-ripple abundance predicts later memory impairment in AD-model mice [14], and soluble tau broadly suppresses cortical activity [15]. Three further substrates shape the failure. Synaptic loss is the structural change that correlates most closely with cognitive decline in AD [16, 17] — the disease has been characterised as a synaptic failure [18] — and hippocampal synapse loss is already measurable at the mild-cognitive-impairment stage [19], providing the substrate for reduced plasticity and replay. GABAergic (notably parvalbumin) interneuron dysfunction disrupts spindle generation and excitation–inhibition balance in cortex, hippocampus and the thalamic reticular nucleus [20, 21]. And degeneration of basal-forebrain cholinergic neurons [22, 23] is implicated in disrupted NREM sleep regulation. Notably, acetylcholine is not a driver of ripple generation: acutely, cholinergic tone *suppresses* hippocampal ripples, which occur precisely when cholinergic tone is at its NREM minimum [24]; cholinergic loss in AD is therefore better understood as a destabiliser of NREM state than as a brake on replay.

Biophysically detailed thalamocortical models have established that SOs and spindles play distinct, complementary roles in spike-timing-dependent plasticity (STDP)-driven consolidation [25, 26], building on the canonical PY–IN–TC–RE network architecture [27] and its neuromodulatory control of sleep stages [28]. Related models show that sleep replay protects memories from catastrophic forgetting [29] and that autonomous hippocampal–neocortical replay supports consolidation [30], and a recent multi-agent formulation casts biological memory as decentralised, adaptive agents [31]. Three questions nevertheless remain open. First, these models treat pathology as a single lesion rather than as a *progressive failure cascade* in which the SO–spindle–ripple machinery degrades step by step as pathology accumulates. Second, we are not aware of a model that dissociates A*β* from tau, showing how two pathologies can produce comparable behavioural deficits through orthogonal electrophysiological signatures. Third, and most consequentially for translation, we are not aware of a model that asks whether a given oscillation-targeting therapy should be expected to help all patients or only a mechanistically defined subgroup.

We address these gaps with a systems-level model pitched at the altitude of neural-mass and mean-field sleep models [32, 33] but coupled to a genuine spiking-plasticity memory module. The model’s design resolves a specific epistemic problem: in many phenomenological models, “coupling” is a latent parameter that is correlated with, but does not cause, consolidation. Here, the *measured* per-event SO–spindle timing alignment, the same quantity read by the modulation index from the simulated field potential [34], directly gates the per-event plasticity. This lets us demonstrate, rather than assume, that consolidation tracks coupling, using a causal control that scrambles timing while holding oscillation power fixed. We then map five neurodegeneration parameters to mechanisms, reproduce the progressive failure cascade, dissociate A*β* from tau, and derive a falsifiable prediction about the specificity of closed-loop slow-oscillation enhancement [35–37].

## Results

### A model in which measured coupling causally gates consolidation

The model comprises five neuronal populations (cortical pyramidal and interneurons; thalamocortical relay and reticular neurons; hippocampal cells) and runs a wake-encoding → N2 →N3 →wake-test cycle. Cortical SOs (~ 0.8 Hz) and thalamic spindles (~ 12 Hz, envelope SD 200 ms) are produced by reduced oscillators; in N3, spindles are recruited near each SO up-state with a degeneration-dependent timing jitter, reproducing the canonical nesting of spindles on the depolarising up-state (Fig 1). Three memory sequences are encoded with graded initial strength (strong, intermediate, weak). During N3, hippocampal ripples replay these sequences; each spindle window opens an STDP gate whose efficacy is the *realized* timing alignment of that spindle to its SO up-state, memories compete to be reactivated, and ordered replays potentiate the winning sequence’s synapses. A 60-trial cued-recall test after sleep reads out trace integrity. Full equations are in Materials and methods. For tractability the spindles are imposed on the SO up-state rather than emerging from a conductance-based reticular–relay loop, and only slow (frontal, ~12 Hz) spindles are represented; we return to these abstractions in the Discussion.

**Figure 1:** Model architecture and healthy baseline. Spike rasters for cortical pyramidal (PY) and inter-(IN) neurons, thalamocortical relay (TC) and reticular (RE) neurons, and hippocampal (HC) cells across the wake–N2–N3–wake cycle, with the simulated local field potential (LFP) and a 6-s N3 zoom showing spindles nested on SO up-states. The network produces a sustained ~0.8 Hz slow oscillation in N3 with spindles riding the up-states, in contrast to the silent fixed point of a precursor conductance-based model.

At healthy settings (100 random seeds), the simulated field potential showed genuine SO–spindle phase–amplitude coupling: the Tort modulation index (MI) in N3 was 0.138. Across ten independent seeds the measured MI (mean 0.137, range 0.131–0.144) exceeded that seed’s own circular-shift surrogate 95th percentile (mean 0.069, range 0.043–0.098) in 10 of 10 cases, confirming the coupling is not a filtering artefact. Consolidation was graded by encoding strength, as expected from competition that favours strong traces [25]: post-sleep recall was 0.978 (95% CI 0.971–0.986; median 0.983, IQR 0.979–0.988) for the strong memory, 0.872 (0.820–0.925; median 0.976, IQR 0.951–0.986) for the intermediate, and 0.488 (0.409–0.567; median 0.530, IQR 0.038–0.934) for the weak memory. The wide interquartile ranges of the weaker traces are not measurement noise but the signature of a winner-take-all competition.

To test whether coupling *causes* consolidation or merely co-varies with it, we scrambled the SO–spindle timing, placing the same number of N3 spindles at random phases instead of on the up-state. Slow-oscillation power was unchanged to four significant figures (875.0 in both conditions), and N3 spindle power was not reduced but slightly *increased* (26.7 → 27.8 a.u., +4.3%, because random placement alters spindle overlap), so the manipulation is conservative with respect to oscillation power. It nonetheless collapsed both the measured coupling (MI 0.138 → 0.007) and memory (grand-mean recall 0.780 → 0.172; Welch *p* = 3.6 × 10^−69^, *d* = 6.43). Because oscillation power was held fixed, the loss of memory is attributable to the loss of coupling *timing* per se (the quantity the plasticity gate reads) rather than to any change in the rhythms’ magnitude. Since the per-event gate multiplies plasticity by the realized alignment, this control establishes that coupling timing is *necessary within the model*, a structural rather than empirical property; it distinguishes the present model from formulations in which coupling and consolidation merely share upstream parameters, but it is not independent biological proof that coupling drives memory.

### Progressive neurodegeneration dismantles the consolidation cascade

We then advanced eight stages of combined pathology, co-varying all five parameters from healthy to severe. Every component of the machinery degraded monotonically (Fig 2): SO–spindle coupling (MI) fell from 0.138 to 0.005, slow-wave (SO) power from 875 to 25 a.u., N2 spindle power from 32.5 to 0.27 a.u., and replay-order fidelity from 1.00 to 0.04. Post-sleep recall fell for all three memories, and the *sleep effect* (Δrecall = post − pre) flipped from net consolidation at early stages to net forgetting at moderate-to-severe stages: the model’s analogue of sleep changing from protective to harmful as the machinery that normally reorganises traces instead lets un-replayed traces fade [29]. The weak memory crossed into net forgetting first and the strong memory last, reproducing differential vulnerability. The phase–amplitude comodulograms show spindle amplitude, concentrated on the SO up-state in health, progressively dispersing with pathology (Fig 4). Across all stages and seeds, per-run MI and per-run mean recall were strongly correlated (Pearson *r* = 0.80, *p <* 10^−100^, *n* = 800; Spearman *r* = 0.82). Grand-mean recall fell from 0.780 to 0.113 between the healthy and most severe stages (Welch *p <* 10^−100^, *d* = 7.07), and the sleep effect for the strong memory crossed from +0.571 to −0.245, changing sign at stage 2. We emphasise the correct interpretation of this association together with the causal control above: the across-condition correlation reflects both shared degeneration and the shared timing mechanism, whereas *within* a fixed disease stage the MI–recall correlation is weak (mean *r* ≈ 0.04), because most across-seed recall variance there arises from the stochastic replay competition rather than from coupling. The causal evidence that coupling drives consolidation is the timing-scramble control, not the across-condition correlation.

**Figure 2:** The neurodegeneration cascade. Mean ± SEM across 100 seeds for SO–spindle coupling (MI), slow-oscillation power, spindle power, per-memory post-sleep recall, the overnight sleep effect (Δrecall), and a mechanistic coupling-efficacy summary, across eight stages of progressive pathology. Coupling, power, and recall fall monotonically; the sleep effect crosses zero into net forgetting, weak memory first. Absolute recall values are set by the cued-recall read-out rather than by the consolidation mechanism (S1 Fig), so only relative changes should be interpreted.

Because consolidation is decided by a finite number of stochastic replay competitions, single-run outcomes are variable; the strong *>* intermediate *>* weak ordering is a cohort-level property. It holds in 52% of seeds at the healthy ceiling (where strong and intermediate both saturate) and rises to 94% under moderate-to-severe pathology as competition separates the traces (Fig 3; Fig 9C). We therefore report recall relative to healthy and treat differential vulnerability as a population effect.

**Figure 3:** Per-seed spread of differential vulnerability. Per-seed post-sleep recall for each memory across the neurodegeneration stages (*N* = 100 seeds per stage; one point per seed, lines are means). The strong *>* intermediate *>* weak ordering is a cohort-level average over a stochastic replay competition rather than a per-run certainty, and mean ± SEM alone would misrepresent these distributions.

**Figure 4:** Loss of phase–amplitude coupling. SO-phase × spindle-amplitude comodulograms (polar histograms of spindle amplitude across SO phase) for healthy, moderate, and severe stages. Healthy sleep shows spindle amplitude concentrated on the SO up-state; this concentration disperses as pathology accumulates.

### Amyloid and tau impair consolidation through orthogonal signatures

We next isolated A*β*-like and tau-like pathology, choosing magnitudes that produce *comparable* behavioural deficits in order to contrast their electrophysiological and replay signatures (the matched-deficit design is a deliberate analytical choice, not a discovered invariant). A*β*-dominant pathology collapsed slow-wave power (SO power 137 versus 875 in health) while leaving hippocampal replay order largely intact (fraction ordered 0.95), reducing strong-memory recall to 0.645 (95% CI 0.601–0.689). Tau-dominant pathology did the opposite: it preserved slow-wave power (SO power 750) but collapsed replay order (fraction ordered 0.65), reducing strong-memory recall to 0.515 (0.476–0.554) (Fig 5). The two deficits were broadly similar in size, though not statistically equivalent (see below), yet arose through orthogonal mechanisms: a low-SWA/intact-replay signature for A*β* and an intact-SWA/degraded-replay signature for tau. Because those two conditions use different pathology magnitudes, we also compared the pathologies at matched knob values. Raising amyloid alone from 0 to 0.9 reduced slow-wave power by 86% (875 → 120) but the modulation index by only 14% (0.138 → 0.119), whereas raising tau alone across the same range reduced slow-wave power by just 20% (875 → 697) yet the modulation index by 62% (0.138 → 0.052). Coupling is thus strongly tau-driven and comparatively amyloid-insensitive, while slow-wave power is the reverse — precisely the pattern reported in humans, where impaired SO– spindle coupling predicts medial-temporal tau while A*β* predicts diminished slow-wave amplitude, the two being statistically dissociable [9, 10].

**Figure 5:** A*β* versus tau double dissociation. Left: per-memory post-sleep recall for healthy, A*β*-only, tau-only, and combined pathology. Right: signature plane of slow-wave (SO) power against replay-order fidelity; A*β* moves the network left (low SWA, intact replay) while tau moves it down (intact SWA, degraded replay), establishing orthogonal axes.

### The double dissociation is a topology-level property, not a calibration artefact

A matched-deficit design invites the objection that the orthogonal signatures depend on the particular pathology magnitudes chosen. We therefore swept the full A*β*× tau plane (6 × 6 grid, 24 seeds per cell) *without* constraining the behavioural deficits to be equal. Across the entire grid, slow-wave power decreased monotonically with amyloid at every level of tau, and replay-order fidelity decreased monotonically with tau at every level of amyloid. The two axes were close to orthogonal: varying tau across its full range changed slow-wave power by only 73 a.u. on average (against a healthy value of 875), whereas varying amyloid across its full range changed replay-order fidelity by only 0.019 on a 0–1 scale. The signature plane is therefore a topology-level property of the model rather than a feature of one calibration point (Fig 6A,B).

**Figure 6:** The A*β*/tau dissociation is a topology-level property. (A) Signature plane over a 6 × 6 A*β*× tau grid (24 seeds per cell), coloured by mean recall; red lines join constant-amyloid rows, blue lines constant-tau columns. Slow-wave power varies almost exclusively along the amyloid axis and replay-order fidelity almost exclusively along the tau axis. (B) Mean recall over the same plane. (C) Resampling all ten disease-slope coefficients within ±50% preserves the dissociation in 40/40 samples. Axes give each pathology’s relative effect on the two readouts: values below 1 on the abscissa mean A*β* reduces slow-wave power more than tau does, and below 1 on the ordinate that tau degrades replay order more than A*β* does, so the lower-left quadrant is the preserved dissociation.

We next asked whether the dissociation depends on the specific coefficient values of the knob-to-mechanism map. Resampling all ten disease-slope coefficients simultaneously from a ±50% uniform range, the dissociation — amyloid depressing slow-wave power more than tau, and tau depressing replay-order fidelity more than amyloid — was preserved in 40 of 40 resamples (100%; Fig 6C).

At *N* = 100 the two conditions are no longer statistically indistinguishable: A*β*-only and tau-only produce comparable but significantly different strong-memory deficits (recall 0.645, 95% CI 0.601–0.689 versus 0.515, 0.476–0.554; Welch *p* = 1.8 × 10^−5^, *d* = 0.62), and equivalence within ±0.10 is not supported (TOST *p* = 0.85). We therefore do **not** claim matched deficits. The dissociation rests instead on the orthogonality of the signatures across the whole plane, which does not require the two deficits to be equal.

### Therapeutic rescue is partial and dissociates coupling from memory

We modelled two interventions drawn from the experimental literature: slow-oscillation enhancement, abstracting closed-loop auditory stimulation and slow-oscillatory transcranial stimulation [35–37], with stimulation timed to the down-to-up transition as in biophysical work [38], implemented as a boost to effective SO amplitude (capped at the healthy value, to avoid a supraphysiological rescue); and GABAergic rescue, abstracting interneuron-targeted restoration of spindle generation [39], implemented as a boost to effective inhibition. Starting from an early–mid-AD profile dominated by the reversible coupling problem, both monotherapies helped and the combination helped most: grand-mean recall rose from 0.152 (no treatment) to 0.172 (SO-enhancement), 0.261 (GABA rescue), and 0.326 (combined). All three arms survived Benjamini–Hochberg correction across the three between-arm comparisons (adjusted *p* = 3.1 × 10^−6^, *d* = 0.68; 5.7 × 10^−33^, *d* = 2.29; and 6.6 × 10^−49^, *d* = 3.41). Only the GABA and combined arms restored a non-negative sleep effect (Δrecall +0.005 and +0.070, versus −0.104 untreated). Rescue was nonetheless partial, attenuating but not abolishing net forgetting in this regime (Fig 7).

**Figure 7:** Therapeutic rescue of a coupling-dominated deficit. Left: grand-mean post-sleep recall per therapy arm, annotated with the N3 modulation index. Right: overnight sleep effect (Δrecall) per arm. SO-enhancement and GABA rescue act through different sub-mechanisms; the MI annotation deliberately illustrates that coupling and memory benefit dissociate across arms. As elsewhere, absolute recall is set by the read-out; the comparison between arms is the interpretable quantity.

A non-trivial dissociation emerged within the therapy arms: SO-enhanceme restored slow-wave power dramatically (from 140 to 809 a.u.) but lifted recall only modestly (*d* = 0.68), whereas GABA rescue left slow-wave power unchanged at 140 yet produced a far larger memory benefit (*d* = 2.29). The modulation index is therefore *not* a clean surrogate for the memory benefit of a therapy, a caution directly relevant to trials that adopt coupling metrics as primary endpoints.

### A falsifiable prediction: slow-oscillation enhancement fails in a tau-dominated deficit

The preceding dissociation implies a specific, testable prediction. We constructed a *tau-dominated* deficit, in which the bottleneck is degraded hippocampal replay rather than weak slow waves, and applied the same therapies. Slow-oscillation enhancement more than doubled slow-wave power (422 → 965 a.u.) yet produced no detectable memory benefit (grandmean recall 0.115 → 0.116; BH-adjusted *p* = 0.62, *d* = 0.07). Because a non-significant test is not by itself evidence of absence, we tested equivalence directly: two one-sided tests supported equivalence within a margin of ±0.05 recall (TOST *p <* 10^−4^; 90% CI on the difference −0.002 to +0.003), and the design had 94% power to detect *d* = 0.5 and 69% power at *d* = 0.35. GABA rescue produced only a small benefit in this regime (0.122; adjusted *p* = 1.8 × 10^−4^, *d* = 0.56), and even combined therapy left the sleep effect firmly negative (Δrecall −0.131 versus −0.141 untreated) (Fig 8). The model’s interpretation is mechanistically transparent: stimulation can re-open and re-time the cortical STDP gate, but cannot repair the corrupted hippocampal replay that the gate is meant to admit. The clinical corollary is that closed-loop SO-enhancement should rescue memory in amyloid/coupling-dominated patients but fail in tau-dominated patients *despite normalising their slow-wave EEG signature*, a directly falsifiable, stratification-relevant prediction.

**Figure 8:** Falsifiable prediction in a tau-dominated deficit. Therapy arms applied to a tau-dominated profile. SO-enhancement restores slow-wave power but does not improve recall, because the deficit lies in replay fidelity, which stimulation cannot repair.

**Figure 9:** What the simulation adds beyond its own mean trajectory. (A) A closed-form prediction combining the potentiation drift budget with the residual un-replayed trace reproduces seed-mean recall across all eight stages (*r* = 0.97, MAE = 0.030); the severe-stage plateau coincides with the residual trace (dotted line), so the mean trajectory is not evidence of emergence. (B) Where competition is active, single-run outcomes are winner-take-all: the healthy weak memory is strongly bimodal (39% of seeds below 0.2, 33% above 0.8; bimodality coefficient 0.72), so the mean describes almost no individual run. (C) The strong ≥ intermediate ≥ weak ordering is probabilistic, rising from 52% of seeds at the healthy ceiling to 94% under pathology, against a chance level of 1/6. *N* = 100 seeds per stage.

### The qualitative results are robust to parameter perturbation

Because the model contains hand-chosen coefficients, we performed a one-at-a-time sensitivity analysis, perturbing each exposed coefficient by ±25% at a mid-stage deficit and measuring the change in grand-mean recall (S1 Fig). The plasticity- and competition-mechanism coefficients (STDP gain, competitive depression, competition softness) each changed recall by less than 0.02, so the dose-response and dissociation results do not depend on their precise values. We additionally perturbed the inline disease-slope coefficients of the knob-to-mechanism map (S1 Fig, right panel): the largest single-coefficient effect on grand-mean recall was 0.004 over ±25%, so the cascade and dissociation do not depend on the specific mapping constants. The dominant sensitivity overall was to the recall-test read-out parameters (the logistic threshold and steepness; S1 Fig, left panel), which calibrate the measurement model rather than the consolidation mechanism; we therefore report recall relative to healthy and verify that orderings, not absolute values, are preserved.

### The simulation adds structure beyond the knob mapping

A fair objection to any phenomenological model is that its averaged behaviour may be an algebraic restatement of its parameter map. We tested this directly, and for the mean the objection is largely correct. An analytic prediction combining a drift budget for potentiation (the product of opportunity rate, ripple availability, nesting probability, replay fidelity, coupling quality and STDP gain; Materials and methods) with the residual un-replayed trace surviving homeostatic downscaling reproduces the seed-averaged recall trajectory closely (Pearson *r* = 0.97, mean absolute error 0.030 in recall units across the eight stages; Fig 9A). In particular the plateau that simulated recall reaches at severe stages (0.113) is not an emergent saturation: it is exactly the residual encoded trace read through the recall test in the absence of any potentiation (analytic value 0.112). We state this plainly because it bounds what the mean dose-response can be claimed to demonstrate.

The simulation’s contribution is therefore distributional, not average. Two features are not captured by any closed form over the means. First, consolidation is genuinely winner-take-all where competition is active: in the healthy condition the weak memory’s recall distribution is strongly bimodal (Sarle’s bimodality coefficient 0.72; 39% of seeds below 0.2 and 33% above 0.8, median 0.53, interquartile range 0.04–0.93), so the reported mean of 0.49 describes a value almost no individual run realises (Fig 9B). This bimodality is a property of the replay competition and is confined to conditions where competition has room to operate; under moderate-to-severe pathology the distributions become unimodal and simply low (stage 3 strong memory: bimodality coefficient 0.39, interquartile range 0.17–0.22). Second, the strong *>* intermediate *>* weak ordering is probabilistic rather than fixed, holding in 52% of seeds at the healthy ceiling and rising to 94% under moderate-to-severe pathology, against a chance level of 17% (Fig 9C). The combination of A*β* and tau, by contrast, was multiplicative (observed combined recall 0.14 versus a multiplicative-null prediction of 0.14) and is not claimed as an emergent interaction.

## Discussion

We have presented a systems-level model of sleep-dependent memory consolidation in which the *measured* SO–spindle timing alignment causally gates synaptic plasticity, and have used it to characterise how progressive neurodegeneration dismantles the consolidation cascade. Four results stand out. First, a timing-scramble control that holds oscillation power fixed abolishes consolidation, establishing, rather than assuming, that coupling drives memory. Second, progressive pathology produces a monotonic collapse of coupling, slow-wave power, spindle power and recall, with the overnight effect flipping from consolidation to net forgetting and weak memories failing first. Third, A*β* and tau impair memory through orthogonal signatures (low-SWA/intact-replay versus intact-SWA/degraded-replay), matching the dissociable human sleep signatures of the two pathologies [9, 10]; this orthogonality is a topology-level property, holding across the whole A*β* × tau plane and surviving ±50% resampling of every mapping coefficient. Fourth, the model predicts that closed-loop slow-oscillation enhancement should help amyloid/coupling-dominated but not tau-dominated patients, even though it normalises slow-wave power in both.

### Relation to prior work

The directional mappings reproduce well-established empirical relationships: A*β*→ reduced SWA → impaired hippocampus-dependent memory [9]; tau → reduced SO–spindle coupling and reduced low-frequency SWA [10, 11]; tau → degraded ripple-associated replay [13, 14]; GABAergic interneuron dysfunction → impaired spindle generation [20, 21]; and SO–spindle coupling → memory across the lifespan [7, 8]. The model complements biophysically detailed thalamocortical simulations [25–28] by trading conductance-level realism for the statistical power and interpretability needed to run an eight-stage ×24-seed disease sweep with formal inference, and by adding what those models do not isolate: a progressive failure cascade, an A*β*/tau double dissociation, and a stratification-relevant therapy prediction. Its abstraction is consistent with neural-mass approaches to NREM rhythms and their response to stimulation [32, 33].

### Clinical implications

The therapy results carry a concrete translational message. Several closed-loop and transcranial stimulation paradigms enhance slow oscillations, spindles, and overnight memory in healthy older adults and in mild cognitive impairment [35–37, 40], and pharmacological and optogenetic restoration of slow waves reduces pathology and improves memory in AD models [39, 41, 42]. Our model suggests these benefits should be conditional on the dominant pathology: where the bottleneck is slow-wave amplitude or coupling (an A*β*-leaning profile), stimulation should help; where the bottleneck is hippocampal replay fidelity (a tau-leaning profile), stimulation may normalise the EEG without rescuing memory. Because A*β* and tau burden are separately measurable in living patients, this prediction implies a biomarker-based stratification for sleep-targeted trials and cautions against adopting coupling metrics as standalone surrogate endpoints.

### Comparison to empirical data and a falsification test

The model’s *relative* effects fall within empirically observed directions and magnitudes: SO–spindle coupling declines with age and medial-temporal tau [7, 10]; slow-wave activity falls with medial-prefrontal amyloid [9]; spindle density and power fall in AD [10, 20]; ripple-associated replay fidelity falls with tau [13, 14]; and the overnight memory benefit declines with both pathologies [9]. We did not fit the model to a dataset, and its absolute units are arbitrary; the results are therefore hypothesis-generating, and the appropriate next step is a direct test of the central prediction. Concretely, in cognitively unimpaired-to-early-AD adults stratified by amyloid and tau PET into amyloid-dominant and tau-dominant groups (≈30 per arm, powering the amyloid-dominant arm for a medium between-group effect of *d* = 0.7 at 80% power — comparable to the smallest therapy effect the model produces in the coupling-dominated regime, *d* = 0.68), a single night of closed-loop slow-oscillation stimulation should improve overnight retention in the amyloid-dominant but not the tau-dominant group, *despite comparable increases in slow-wave power in both*. Slow-wave restoration without memory benefit in the tau-dominant arm would corroborate the model; a uniform benefit across strata would falsify its central claim. Existing sleep-stimulation cohorts in mild cognitive impairment [36, 40], if retrospectively stratified by amyloid and tau biomarkers, could provide a first test, in particular by asking whether the responders and non-responders in [40] differ in tau burden. Table 1 maps each headline model output onto the corresponding human observation. Two points of interpretation matter: the model’s *absolute* values are not comparable to published ones (the Tort MI depends on bin count, band edges and epoch length, and our amplitudes are in arbitrary units), so only direction and relative change are claimed; and one observation speaks directly to the central prediction — in amnestic MCI, acoustic slow-oscillation stimulation raised slow-wave activity by more than 10% over sham yet improved recall in only five of nine patients [40]. Our model supplies a testable account of that responder heterogeneity: the non-responders may be the tau-dominant subgroup.

**Table 1:** Model outputs against human observations. Absolute values are not comparable across studies (see text); direction and relative change are.

| Model output | Model (healthy → severe) | Human observation | Source | Agreement |
| --- | --- | --- | --- | --- |
| SO-spindle coupling (MI) | 0.138 → 0.005 | Coupling quality predicts overnight retention; mPFC atrophy causes temporal dispersion of coupling and forgetting | [7] | Direction only (absolute MI not comparable) |
| Which pathology drives coupling loss | $\tau \gg A\beta$ | Impaired coupling predicts medial-temporal $\tau$; $A\beta$ instead predicts reduced <1 Hz SWA, statistically dissociable | [10] | Reproduces reported dissociation |
| Slow-wave (SO) power | 875 → 25 a.u. | mPFC $A\beta$ correlates with impaired NREM SWA generation; SWA inversely related to $\tau$ at 1–2 Hz | [9, 11] | Direction; units arbitrary |
| Spindle power | 32.5 → 0.27 a.u. | Spindle measures reduced in AD | [10, 20] | Direction |
| Replay-order fidelity | 1.00 → 0.04 | Tau disrupts SWR spike dynamics; reduced SWR abundance predicts later memory loss | [13, 14] | Direction |
| Overnight memory effect | +0.571 → -0.245 (sign change) | Reduced SWA impairs overnight consolidation; coupling dispersion produces forgetting | [7, 9] | Direction |
| SWA restored without memory gain | tau-dominated: SO power 422 → 965, recall unchanged (0.115 → 0.116) | Acoustic stimulation in aMCI raised SWA >10% but improved recall in only 5/9 patients | [40] | Consistent with responder heterogeneity |
| Coupling restored with memory gain | combined therapy raises coupling and recall | so-tDCS in MCI increased SO/spindle power and synchronisation and improved memory | [36] | Direction |

### Limitations

This is a phenomenological model and makes no conductance-level biophysical claim; its numbers are relative and qualitative. Four caveats deserve emphasis. (i) The *seed-averaged* dose-response curves are interpretable, near-closed-form consequences of the parameter-to-mechanism mapping; the model’s added value lies in the variance structure (stochastic competition, differential vulnerability, per-event gating) and in the causal control, not in the mean trend, which could be written analytically. (ii) Within a fixed disease stage the MI–recall correlation is weak, because the replay competition dominates across-seed variance; the causal evidence for the coupling-to-memory link is the timing-scramble control, not the correlation. (iii) We do not claim matched A*β*/tau deficits: at *N* = 100 the two conditions differ significantly (*p* = 1.8 × 10^−5^, *d* = 0.62) and equivalence within ±0.10 is not supported. The dissociation claim rests instead on the orthogonality of the signature axes across the whole A*β*× tau plane and on its survival of ±50% coefficient resampling. (iv) Absolute units are arbitrary (we therefore report relative recall); the sensitivity analysis now spans the read-out, plasticity, and inline mapping coefficients, with the recall read-out the dominant lever. We also model acetylcholine as a NREM-state-stability factor rather than a ripple scheduler, consistent with evidence that acute cholinergic tone suppresses ripples [24]; chronic cholinergic degeneration is represented through its destabilising effect on NREM maintenance [22, 23]. The thalamic reticular–relay spindle loop is represented at the mean-field level rather than as a conductance-based reciprocal circuit; coupling the present plasticity module to such a network [27, 28] is a natural extension. The GABA-to-SWA arm is the weaker-cited of the two GABA effects and is better read as excitation–inhibition degradation than as direct amplitude scaling.

Three further scope limitations deserve explicit treatment. First, **only slow, frontally dominant spindles (**~**12 Hz) are represented**. Fast (~13–16 Hz), centro-parietal spindles couple to the SO up-state with a distinct phase preference and are the subtype most closely tied to hippocampus-dependent consolidation [4, 5]. Because tau acts in our model by dispersing spindle-to-SO timing, adding a second spindle population with its own phase preference should *sharpen* rather than blunt the dissociation: fast-spindle coupling would be the more tau-sensitive readout, while slow-spindle amplitude would track the A*β*/GABA amplitude axis. This is a concrete, testable consequence of the present architecture rather than a defect of it. Second, **REM sleep is omitted**. Since tau pathology also degrades REM-associated theta activity and reactivation in rodent models, and REM is implicated in stabilising and integrating newly consolidated traces, this omission most plausibly causes the model to *underestimate* the tau-associated deficit, while leaving the A*β*-associated deficit — which acts through NREM slow-wave amplitude — comparatively unaffected. The direction of this bias is therefore conservative with respect to our central dissociation claim. Third, **the cholinergic knob acts only as an NREM-state-stability factor**. This is well supported for ripple scheduling, since acute cholinergic tone suppresses ripples [24], but it simplifies the wider cholinergic contribution: muscarinic signalling also modulates cortical excitability and slow-wave generation, so a fuller treatment would allow acetylcholine to act on SO generation directly as well as on state maintenance. Two further simplifications should be noted. The derived rate of roughly 3.3 ripples per spindle window (an implied ~2.5 ripple events per second) sits at the upper end of, or slightly above, reported NREM sharp-wave-ripple rates; the scaling constant that sets it is declared as freely tuned (S1 Table) and the qualitative results do not depend on it (S1 Fig). The spindle carrier is also held at a fixed 12 Hz, whereas real spindles show within-event frequency drift. Circadian modulation and glymphatic clearance are likewise outside the model’s scope. A caveat applies to the A*β* arm specifically: in APP/PS1 mice, A*β* alone reduces cortical slow-oscillation power *and* degrades the coordination of hippocampal sharp-wave ripples with both the slow oscillation and thalamocortical spindles [43]. Our mapping, in which A*β* leaves replay order essentially intact, is therefore an idealisation that sharpens the contrast with tau; a more graded A*β* effect on replay coordination would blunt, though not reverse, the dissociation.

### Future directions

The model makes several testable predictions beyond the therapy result: that the temporal *precision* of SO–spindle coupling, not slow-wave power, should track memory in tau-dominant patients; that interventions restoring spindle generation (e.g., interneuron-targeted approaches) should benefit memory more than slow-wave amplitude alone in coupling-limited regimes; and that overnight forgetting, rather than merely reduced consolidation, should emerge once the machinery passes a threshold. Validating the therapy-specificity prediction against A*β*/tau-stratified human stimulation data is the natural empirical next step.

## Materials and methods

### Model overview and rhythms

The model represents five populations (120 cortical pyramidal, 30 cortical inter-, 40 thalamocortical relay, 40 reticular, 20 hippocampal units) and simulates a wake-encode (2 s) → N2 (10 s) → N3 (24 s) → wake-test (2 s) cycle at 1 ms resolution. Spiking rasters are generated for visualisation; the consolidation mechanism operates on sequence-synapse weights and is read out behaviourally (below). The slow oscillation is a phase oscillator advancing at *f*_SO_ = 0.8 Hz with cycle-to-cycle frequency jitter, with waveform

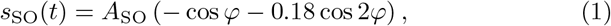

giving a sharpened up-state at *φ* = *π*. In N3 the SO is expressed at full amplitude *A*_SO_; in N2 only a residual SO is present. Spindles are 12 Hz carriers under a Gaussian envelope (SD 200 ms; full-width at half-maximum ≈470 ms, yielding detected durations of ~0.5–1.2 s, within the canonical range). In N3, a spindle is recruited on each SO up-state with probability *p*_SO-sp_; its centre is placed at the up-peak plus a nominal phase offset plus Gaussian timing jitter, so that each spindle carries a *realized* phase offset Δ*φ* relative to its up-state. In N2, spindles occur as a Poisson process. The simulated LFP is the sum of the SO, spindle, and pink-ish background-noise components.

### Neurodegeneration parameter-to-mechanism mapping

Five parameters in [0, 1] (amyloid *a*, tau *t*) or [~ 0.1, 1] (synaptic density *s*, GABAergic efficiency *g*, cholinergic tone *c*) map monotonically to mechanism variables (Table 2). For example, the effective SO amplitude is

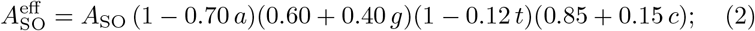

spindle amplitude and occurrence scale with effective inhibition and decline modestly with *a* and *t*; spindle-to-SO jitter increases with *t* and with reduced SO amplitude; replay-order fidelity declines with *t* and *s*; and ripple supply is substrate-driven (*s, t*) and scaled by NREM-state stability (a function of *c*), not positively gated by acetylcholine.

**Table 2:** Neurodegeneration parameter-to-mechanism mapping. ↓/↑ denote decrease/increase with worsening pathology.

| Parameter | Primary mechanistic effect | Key references |
| --- | --- | --- |
| amyloid ($a$) | ↓ SO amplitude / slow-wave activity; weak up-states recruit spindles at a sloppier phase | [9] |
| tau ($t$) | ↓ replay order fidelity and per-link timing;<br>↑ spindle-to-SO jitter (decoupling); modest ↓ spindle/SWA | [10, 13, 14] |
| synaptic density ($s$) | ↓ STDP gain, ripple supply, recall transmission | [16–19] |
| GABA ($g$) | ↓ spindle amplitude/occurrence (RE/interneuron); ↓ SO amplitude (E/I) | [20, 39] |
| ACh ($c$) | ↓ NREM-state stability (SO/spindle maintenance, coordinated-window availability); *not* a ripple scheduler | [22–24] |

### Replay, coupling-gated plasticity, and recall

Three sequences (length 8) are encoded with graded initial weights proportional to encoding strength (1.0, 0.72, 0.5). During N3, each spindle window admits a Poisson number of nested ripples (a function of ripple rate). For each ripple: (i) a ripple is delivered with probability equal to ripple availability; (ii) it is nested in the usable window with probability *coupled-window*; (iii) memories compete for reactivation, with

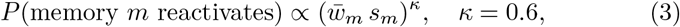

so strong traces are favoured but weak traces occasionally win [25]; (iv) replay is correctly ordered with probability equal to replay fidelity. The per-event coupling gate is the realized timing alignment of the host spindle,

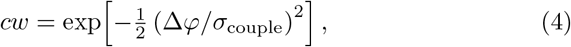

where *σ*_couple_ is half the up-state width; this is the same realized timing the modulation index reads from the LFP. For an ordered, nested replay, each sequence link potentiates only if its replayed spike falls within the ~25 ms STDP window [44] (probability declining with tau-driven replay jitter), with increment *g*_STDP_ · *cw* · *a*_amp_, where *a*_amp_ is the spindle’s amplitude factor; competing memories undergo small competitive depression. Homeostatic down-scaling is applied across N3 so that un-replayed traces fade. Weights are bounded. After sleep, a cued-recall test runs 60 stochastic trials per memory; each sequence link transmits with probability

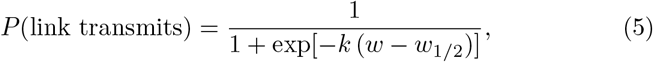

and trace integrity is the fraction of links transmitting. The sleep effect (Δrecall) is post-sleep minus pre-sleep recall. The causal control sets a coupling-shuffle flag, placing N3 spindles at random phases so that SO and spindle power are preserved but the per-event gate and MI collapse.

### Signal analysis

The modulation index is computed by the Tort method [34] from the SO phase (0.5–1.5 Hz) and spindle amplitude (10–16 Hz) of the simulated LFP using 18 phase bins. A circular time-shift surrogate (200 iterations) provides a coupling null; the real MI is reported against the surrogate mean and 95th percentile. Band power is computed by Welch’s method (8 s segments). Spindle and SO detectors are retained only as labelled diagnostics; because count-based detector outputs (spindle density, SO frequency) are threshold/zero-crossing artefacts that do not track the generative parameters, the headline rhythm metrics are band power (SO power, spindle power) and the MI.

### Statistics and reproducibility

Unless stated otherwise, conditions were run with 100 random seeds; figures show mean ± SEM and the text reports 95% confidence intervals (t-distribution) together with medians and interquartile ranges, since several recall distributions are strongly bimodal. Between-condition comparisons use Welch’s *t*-test with Cohen’s *d* (Mann–Whitney *U* as a non-parametric check). Within each experimental block, all between-arm comparisons are corrected for multiple testing by the Benjamini–Hochberg procedure and adjusted *p*-values are reported. Where a null result is interpreted substantively, it is supported by two one-sided tests of equivalence (TOST) with an explicit margin and by the achieved power at the effect sizes of interest, rather than by a non-significant test alone. The MI–recall relationship is reported as a Pearson and Spearman correlation pooled across stages and seeds, and as the mean within-condition correlation. Robustness is assessed by one-at-a-time ±25% perturbation of exposed coefficients. The model is fully deterministic given a seed; an independent execution on a separate platform reproduced all reported numbers bit-for-bit.

## Data and code availability

All simulation and analysis code needed to reproduce every figure and numerical result in this paper is openly available at Zenodo: https://doi.org/10.5281/zenodo.22864939 (concept DOI, which resolves to the most recent version). The deposit contains the model library (`sleep_consolidation_model.py`). a single driver script (`run_analysis.py`) that reproduces the complete experiment matrix in approximately 20 minutes on one CPU core, and documentation mapping each output file to the figure, table or reported value it generates. The code requires only Python 3.9 or later with NumPy, SciPy and Matplotlib. The model is deterministic given a random seed; all seeds used for the reported results are documented in the deposit. This study uses no empirical dataset, so no data are deposited or redistributed.

## Ethics declaration

This study is a computational simulation and involved no human participants, animals, or personal data; no ethical approval was required.

## Author contributions

**B.M**.: Conceptualization, Methodology, Investigation, Writing – original draft, Writing – review & editing. **A.S**.: Conceptualization, Methodology, Software, Formal analysis, Visualization, Writing – review & editing. Z.M.: Supervision, Investigation, Writing – review & editing. All authors reviewed the manuscript

## Supporting information

**S1 Fig. Parameter sensitivity (tornado)**. Change in grand-mean recall under ±25% one-at-a-time perturbation at a mid-stage deficit. Left: Config coefficients (plasticity/competition/read-out); the recall read-out parameters are the largest lever (a measurement-model, not mechanism, sensitivity). Right: inline disease-slope coefficients of the knob-to-mechanism map; the largest single-coefficient effect is 0.004, so the cascade and dissociation do not depend on the mapping constants.

**S1 Table. Provenance of every model coefficient**. Each coefficient is declared as [E] empirically constrained in direction and approximate magnitude, [O] an order-of-magnitude estimate anchored to a physiological quantity, or [F] freely tuned (chosen to place the model in a workable dynamic range; no empirical claim). No disease-slope coefficient magnitude is fitted to data; the empirical constraint is on the sign and relative ordering of the dependencies. Disease slopes: *a* →SO amplitude 0.70 [E dir./F mag.] [9]; *t* → SO amplitude 0.12 [E dir., deliberately ≪ the amyloid slope] [10, 11]; *a* → spindle amplitude 0.20 [F]; *t* → spindle amplitude 0.18 [E dir./F mag.] [10]; *a* → *P* (spindle on up-state) 0.25 [E dir./F mag.] [9]; *t* → replay-order fidelity 0.85 [E dir./F mag.] [13,14]; *t* → spindle-to-SO jitter 180 ms [O; ≈14% of the 1250 ms SO cycle] [7, 10]; *t* → ripple supply 0.30, *t* → nesting probability 0.25, *t* → ripple availability 0.25 [all E dir./F mag.] [13, 14]. Rhythm constants: SO 0.8 Hz [E] [1]; spindle 12 Hz [E] [5]; spindle envelope SD 200 ms [E, giving detected durations ~0.5–1.2 s]; healthy coupling jitter 25 ms [O]; ripple rate 1.9 s^−1^ [O]; SO and spindle amplitudes 30 and 14 a.u. [F, only ratios interpreted]. Plasticity and read-out: STDP gain 0.22, competitive depression 0.020, competition softness 0.60, homeostatic decay 0.018 [all F, model-internal]; recall logistic steepness 2.5, half-point 1.35, 60 trials [F, measurement model and the dominant sensitivity, hence recall is reported relative to healthy]; simulated N3 duration 24 s [F, compressed for tractability, not a physiological night]. **Mechanism versus reporting:** the derived quantities entering the plasticity path are `so_amp, spindle_amp, spindle_jitter_ms, so_spindle_prob, n2_spindle_rate, ripples_per_spindle, replay_fidelity, replay_jitter_ms, coupled_win couple_sigma_ms, stdp_gain` and `ripple_availability`. Two are reported but do not gate plasticity: `ripple_rate` (acting only through `ripples_per_spindle`) and `coupling_efficacy` (a plotted summary); they are listed for transparency, not as additional mechanisms.

## Acknowledgments

Not applicable.

## References

[1] Diekelmann S, Born J. The memory function of sleep. Nature Reviews Neuroscience. 2010 Jan;11(2):114–126. Available from: http://dx.doi.org/10.1038/nrn2762. doi:10.1038/nrn2762.

[2] Klinzing JG, Niethard N, Born J. Mechanisms of systems memory consolidation during sleep. Nature Neuroscience. 2019 Aug;22(10):1598–1610. Available from: http://dx.doi.org/10.1038/s41593-019-0467-3. doi:10.1038/s41593-019-0467-3.

[3] McClelland JL, McNaughton BL, O’Reilly RC. Why there are complementary learning systems in the hippocampus and neocortex: Insights from the successes and failures of connectionist models of learning and memory. Psychological Review. 1995;102(3):419–457. Available from: http://dx.doi.org/10.1037/0033-295X.102.3.419. doi:10.1037/0033-295x.102.3.419.

[4] Staresina BP, Bergmann TO, Bonnefond M, van der Meij R, Jensen O, Deuker L, et al. Hierarchical nesting of slow oscillations, spindles and ripples in the human hippocampus during sleep. Nature Neuroscience. 2015;18(11):1679–1686. Available from: http://dx.doi.org/10.1038/nn.4119. doi:10.1038/nn.4119.

[5] Mölle M, Marshall L, Gais S, Born J. Grouping of Spindle Activity during Slow Oscillations in Human Non-Rapid Eye Movement Sleep. The Journal of Neuroscience. 2002 Dec;22(24):10941–10947. Available from: http://dx.doi.org/10.1523/JNEUROSCI.22-24-10941.2002. doi:10.1523/jneurosci.22-24-10941.2002.

[6] Latchoumane CFV, Ngo HVV, Born J, Shin HS. Thalamic Spindles Promote Memory Formation during Sleep through Triple Phase-Locking of Cortical, Thalamic, and Hippocampal Rhythms. Neuron. 2017;95(2):424–435.e6. Available from: http://dx.doi.org/10.1016/j.neuron.2017.06.025. doi:10.1016/j.neuron.2017.06.025.

[7] Helfrich RF, Mander BA, Jagust WJ, Knight RT, Walker MP. Old Brains Come Uncoupled in Sleep: Slow Wave-Spindle Synchrony, Brain Atrophy, and Forgetting. Neuron. 2018 Jan;97(1):221–230.e4. Available from: http://dx.doi.org/10.1016/j.neuron.2017.11.020. doi:10.1016/j.neuron.2017.11.020.

[8] Muehlroth BE, Sander MC, Fandakova Y, Grandy TH, Rasch B, Shing YL, et al. Precise Slow Oscillation–Spindle Coupling Promotes Memory Consolidation in Younger and Older Adults. Scientific Reports. 2019 Feb;9(1). Available from: http://dx.doi.org/10.1038/s41598-018-36557-z. doi:10.1038/s41598-018-36557-z.

[9] Mander BA, Marks SM, Vogel JW, Rao V, Lu B, Saletin JM, et al. β-amyloid disrupts human NREM slow waves and related hippocampus-dependent memory consolidation. Nature Neuroscience. 2015;18(7):1051–1057. Available from: http://dx.doi.org/10.1038/nn.4035. doi:10.1038/nn.4035.

[10] Winer JR, Mander BA, Helfrich RF, Maass A, Harrison TM, Baker SL, et al. Sleep as a Potential Biomarker of Tau and β-Amyloid Burden in the Human Brain. The Journal of Neuroscience. 2019;39(32):6315–6324. Available from: http://dx.doi.org/10.1523/JNEUROSCI.0503-19.2019. doi:10.1523/jneurosci.0503-19.2019.

[11] Lucey BP, McCullough A, Landsness EC, Toedebusch CD, McLeland JS, Zaza AM, et al. Reduced non–rapid eye movement sleep is associated with tau pathology in early Alzheimer’s disease. Science Translational Medicine. 2019 Jan;11(474). Available from: http://dx.doi.org/10.1126/scitranslmed.aau6550. doi:10.1126/scitranslmed.aau6550.

[12] Menkes-Caspi N, Yamin H, Kellner V, Spires-Jones T, Cohen D, Stern E. Pathological Tau Disrupts Ongoing Network Activity. Neuron. 2015 Mar;85(5):959–966. Available from: http://dx.doi.org/10.1016/j.neuron.2015.01.025. doi:10.1016/j.neuron.2015.01.025.

[13] Witton J, Staniaszek LE, Bartsch U, Randall AD, Jones MW, Brown JT. Disrupted hippocampal sharp-wave ripple-associated spike dynamics in a transgenic mouse model of dementia. The Journal of Physiology. 2015 Jan;594(16):4615–4630. Available from: http://dx.doi.org/10.1113/jphysiol.2014.282889. doi:10.1113/jphysiol.2014.282889.

[14] Jones EA, Gillespie AK, Yoon SY, Frank LM, Huang Y. Early Hippocampal Sharp-Wave Ripple Deficits Predict Later Learning and Memory Impairments in an Alzheimer’s Disease Mouse Model. Cell Reports. 2019 Nov;29(8):2123–2133.e4. Available from: http://dx.doi.org/10.1016/j.celrep.2019.10.056. doi:10.1016/j.celrep.2019.10.056.

[15] Busche MA, Wegmann S, Dujardin S, Commins C, Schiantarelli J, Klickstein N, et al. Tau impairs neural circuits, dominating amyloid-β effects, in Alzheimer models in vivo. Nature Neuroscience. 2018 Dec;22(1):57–64. Available from: http://dx.doi.org/10.1038/s41593-018-0289-8. doi:10.1038/s41593-018-0289-8.

[16] Terry RD, Masliah E, Salmon DP, Butters N, DeTeresa R, Hill R, et al. Physical basis of cognitive alterations in alzheimer’s disease: Synapse loss is the major correlate of cognitive impairment. Annals of Neurology. 1991 Oct;30(4):572–580. Available from: http://dx.doi.org/10.1002/ana.410300410. doi:10.1002/ana.410300410.

[17] DeKosky ST, Scheff SW. Synapse loss in frontal cortex biopsies in Alzheimer’s disease: Correlation with cognitive severity. Annals of Neurology. 1990 May;27(5):457–464. Available from: http://dx.doi.org/10.1002/ana.410270502. doi:10.1002/ana.410270502.

[18] Selkoe DJ. Alzheimer’s Disease Is a Synaptic Failure. Science. 2002 Oct;298(5594):789–791. Available from: http://dx.doi.org/10.1126/science.1074069. doi:10.1126/science.1074069.

[19] Scheff SW, Price DA, Schmitt FA, Mufson EJ. Hippocampal synaptic loss in early Alzheimer’s disease and mild cognitive impairment. Neurobiology of Aging. 2006 Oct;27(10):1372–1384. Available from: http://dx.doi.org/10.1016/j.neurobiolaging.2005.09.012. doi:10.1016/j.neurobiolaging.2005.09.012.

[20] Katsuki F, Gerashchenko D, Brown RE. Alterations of sleep oscillations in Alzheimer’s disease: A potential role for GABAergic neurons in the cortex, hippocampus, and thalamus. Brain Research Bulletin. 2022;187:181–198. Available from: http://dx.doi.org/10.1016/j.brainresbull.2022.07.002. doi:10.1016/j.brainresbull.2022.07.002.

[21] Hijazi S, Smit AB, van Kesteren RE. Fast-spiking parvalbumin-positive interneurons in brain physiology and Alzheimer’s disease. Molecular Psychiatry. 2023;28(12):4954–4967. Available from: http://dx.doi.org/10.1038/s41380-023-02168-y. doi:10.1038/s41380-023-02168-y.

[22] Whitehouse P, Price D, Struble R, Clark A, Coyle J, DeLong MR. Alzheimer’s Disease and Senile Dementia: Loss of Neurons in the Basal Forebrain. Science. 1982 Mar;215(4537):1237–1239. Available from: http://dx.doi.org/10.1126/science.7058341. doi:10.1126/science.7058341.

[23] Mesulam MCholinergic circuitry of the human nucleus basalis and its fate in Alzheimer’s disease. Journal of Comparative Neurology. 2013 Oct;521(18):4124–4144. Available from: http://dx.doi.org/10.1002/cne.23415. doi:10.1002/cne.23415.

[24] Zhang Y, Cao L, Varga V, Jing M, Karadas M, Li Y, et al. Cholinergic suppression of hippocampal sharp-wave ripples impairs working memory. Proceedings of the National Academy of Sciences. 2021 Apr;118(15). Available from: http://dx.doi.org/10.1073/pnas.2016432118. doi:10.1073/pnas.2016432118.

[25] Wei Y, Krishnan GP, Komarov M, Bazhenov M. Differential roles of sleep spindles and sleep slow oscillations in memory consolidation. PLOS Computational Biology. 2018;14(7):e1006322. Available from: http://dx.doi.org/10.1371/journal.pcbi.1006322. doi:10.1371/journal.pcbi.1006322.

[26] Wei Y, Krishnan GP, Bazhenov M. Synaptic Mechanisms of Memory Consolidation during Sleep Slow Oscillations. The Journal of Neuroscience. 2016 Apr;36(15):4231–4247. Available from: http://dx.doi.org/10.1523/JNEUROSCI.3648-15.2016. doi:10.1523/jneurosci.3648-15.2016.

[27] Bazhenov M, Timofeev I, Steriade M, Sejnowski TJ. Model of Thalamocortical Slow-Wave Sleep Oscillations and Transitions to Activated States. The Journal of Neuroscience. 2002 Oct;22(19):8691–8704. Available from: http://dx.doi.org/10.1523/JNEUROSCI.22-19-08691.2002. doi:10.1523/jneurosci.22-19-08691.2002.

[28] Krishnan GP, Chauvette S, Shamie I, Soltani S, Timofeev I, Cash SS, et al. Cellular and neurochemical basis of sleep stages in the thalamocortical network. eLife. 2016 Nov;5. Available from: http://dx.doi.org/10.7554/eLife.18607. doi:10.7554/elife.18607.

[29] González OC, Sokolov Y, Krishnan GP, Delanois JE, Bazhenov M. Can sleep protect memories from catastrophic forgetting? eLife. 2020 Aug;9. Available from: http://dx.doi.org/10.7554/eLife.51005. doi:10.7554/elife.51005.

[30] Singh D, Norman KA, Schapiro AC. A model of autonomous interactions between hippocampus and neocortex driving sleep-dependent memory consolidation. Proceedings of the National Academy of Sciences. 2022 Oct;119(44). Available from: http://dx.doi.org/10.1073/pnas.2123432119. doi:10.1073/pnas.2123432119.

[31] Wei H, Feng C, Li F. Modeling biological memory network by an autonomous and adaptive multi-agent system. Brain Informatics. 2024;11(1). Available from: http://dx.doi.org/10.1186/s40708-024-00237-8. doi:10.1186/s40708-024-00237-8.

[32] Schellenberger Costa M, Weigenand A, Ngo HVV, Marshall L, Born J, Martinetz T, et al. A Thalamocortical Neural Mass Model of the EEG during NREM Sleep and Its Response to Auditory Stimulation. PLOS Computational Biology. 2016;12(9):e1005022. Available from: http://dx.doi.org/10.1371/journal.pcbi.1005022. doi:10.1371/journal.pcbi.1005022.

[33] Reato D, Gasca F, Datta A, Bikson M, Marshall L, Parra LC. Transcranial Electrical Stimulation Accelerates Human Sleep Homeostasis. PLoS Computational Biology. 2013 Feb;9(2):e1002898. Available from: http://dx.doi.org/10.1371/journal.pcbi.1002898. doi:10.1371/journal.pcbi.1002898.

[34] Tort ABL, Komorowski R, Eichenbaum H, Kopell N. Measuring Phase-Amplitude Coupling Between Neuronal Oscillations of Different Frequencies. Journal of Neurophysiology. 2010 Aug;104(2):1195–1210. Available from: http://dx.doi.org/10.1152/jn.00106.2010. doi:10.1152/jn.00106.2010.

[35] Ngo HV, Martinetz T, Born J, Mölle M. Auditory Closed-Loop Stimulation of the Sleep Slow Oscillation Enhances Memory. Neuron. 2013 May;78(3):545–553. Available from: http://dx.doi.org/10.1016/j.neuron.2013.03.006. doi:10.1016/j.neuron.2013.03.006.

[36] Ladenbauer J, Ladenbauer J, Külzow N, de Boor R, Avramova E, Grittner U, et al. Promoting Sleep Oscillations and Their Functional Coupling by Transcranial Stimulation Enhances Memory Consolidation in Mild Cognitive Impairment. The Journal of Neuroscience. 2017;37(30):7111–7124. Available from: http://dx.doi.org/10.1523/JNEUROSCI.0260-17.2017. doi:10.1523/jneurosci.0260-17.2017.

[37] Marshall L, Helgadóttir H, Mölle M, Born J. Boosting slow oscillations during sleep potentiates memory. Nature. 2006 Nov;444(7119):610–613. Available from: http://dx.doi.org/10.1038/nature05278. doi:10.1038/nature05278.

[38] Wei Y, Krishnan GP, Marshall L, Martinetz T, Bazhenov M. Stimulation Augments Spike Sequence Replay and Memory Consolidation during Slow-Wave Sleep. The Journal of Neuroscience. 2019 Dec;40(4):811–824. Available from: http://dx.doi.org/10.1523/JNEUROSCI.1427-19.2019. doi:10.1523/jneurosci.1427-19.2019.

[39] Zhao Q, Maci M, Miller MR, Zhou H, Zhang F, Algamal M, et al. Sleep restoration by optogenetic targeting of GABAergic neurons reprograms microglia and ameliorates pathological phenotypes in an Alzheimer’s disease model. Molecular Neurodegeneration. 2023 Dec;18(1). Available from: http://dx.doi.org/10.1186/s13024-023-00682-9. doi:10.1186/s13024-023-00682-9.

[40] Papalambros NA, Weintraub S, Chen T, Grimaldi D, Santostasi G, Paller KA, et al. Acoustic enhancement of sleep slow oscillations in mild cognitive impairment. Annals of Clinical and Translational Neurology. 2019;6(7):1191–1201. Available from: http://dx.doi.org/10.1002/acn3.796. doi:10.1002/acn3.796.

[41] Kastanenka KV, Hou SS, Shakerdge N, Logan R, Feng D, Wegmann S, et al. Optogenetic Restoration of Disrupted Slow Oscillations Halts Amyloid Deposition and Restores Calcium Homeostasis in an Animal Model of Alzheimer’s Disease. PLOS ONE. 2017 Jan;12(1):e0170275. Available from: http://dx.doi.org/10.1371/journal.pone.0170275. doi:10.1371/journal.pone.0170275.

[42] Kollarik S, Bimbiryte D, Sethi A, Dias I, Moreira CG, Noain D. Pharmacological enhancement of slow-wave activity at an early disease stage improves cognition and reduces amyloid pathology in a mouse model of Alzheimer’s disease. Frontiers in Aging Neuroscience. 2025 Jan;16. Available from: http://dx.doi.org/10.3389/fnagi.2024.1519225. doi:10.3389/fnagi.2024.1519225.

[43] Zhou H, Li H, Gowravaram N, Quan M, Kausar N, Gomperts SN. Disruption of hippocampal neuronal circuit function depends upon behavioral state in the APP/PS1 mouse model of Alzheimer’s disease. Scientific Reports. 2022 Dec;12(1). Available from: http://dx.doi.org/10.1038/s41598-022-25364-2. doi:10.1038/s41598-022-25364-2.

[44] Dickey CW, Sargsyan A, Madsen JR, Eskandar EN, Cash SS, Halgren E. Travelling spindles create necessary conditions for spiketiming-dependent plasticity in humans. Nature Communications. 2021 Feb;12(1). Available from: http://dx.doi.org/10.1038/s41467-021-21298-x. doi:10.1038/s41467-021-21298-x.

